# Competition among host-specific lineages of *Poecilochirus carabi* mites influences the extent of co-adaptation with their *Nicrophorus vespilloides* burying beetle hosts

**DOI:** 10.1101/641936

**Authors:** Syuan-Jyun Sun, Rebecca M. Kilner

**Author notes:** **Corresponding author:** Syuan-Jyun Sun.

## Abstract

Reciprocal selection between symbiotic organisms and their hosts can generate variations in local adaptation between them. Symbionts often form species complexes with lineages partially adapted to various hosts. However, it is unclear how interactions among these lineages influences geographic variation in host-symbiont local adaptation. We addressed this shortcoming with experiments on burying beetles *Nicrophorus vespilloides* and their specialist phoretic mite *Poecilochirus carabi* in two adjacent woodlands. Burying beetles transport these mites to vertebrate carrion upon which they both reproduce. *P. carabi* appears to be a species complex, with distinct lineages that specialise on breeding alongside different *Nicrophorus* species. We found that in one wood (Gamlingay Woods), *N. vespilloides* carries a mixture of mite lineages, with each lineage corresponding to one of the four *Nicrophorus* species that inhabits this wood. However, two burying beetle species coexist in neighbouring Waresley Woods and here *N. vespilloides* predominantly carries the mite lineage that favours *N. vespilloides*. Mite lineage mixing alters the degree of local adaptation for both *N. vespilloides* and the *P. carabi* mites, affecting reproductive success variably across different woodlands. In Gamlingay, mite lineage mixing reduced *N. vespilloides* reproductive success, while experimentally purifying mites lineage enhanced it. The near pure lineage of *vespilloides* mites negligibly impacted Waresley *N. vespilloides*. Mite reproductive success varied with host specificity: Gamlingay mites had greatest reproductive success on Gamlingay beetles, and performed less well with Waresley beetles. By contrast, Waresley mites had consistent reproductive success, regardless of beetle’s woodland of origin. We conclude that there is some evidence that *N. vespilloides* and its specific mite lineage have co-adapted. However, neither *N. vespilloides* nor its mite lineage adapted to breed alongside other mite lineages. This, we suggest, causes variation between Waresley and Gaminglay Woods in the extent of local adaptation between *N. vespilloides* beetles and their *P. carabi* mites.

## Introduction

Ever since Darwin (1859), evidence has been gathering that natural selection causes populations to adapt in different ways to their different local environments. More recent work suggests that adaptation can happen on very local scales, even when there is still gene flow between populations (e.g. Sun et al., 2020; J. N. Thompson, 2013). Nevertheless, the extent of local adaptation varies across landscapes, between populations of the same species (e.g. J. N. Thompson, 1999, 2005). This is especially true for adaptations that arise from co-evolutionary interactions between symbiotic species (by which we mean ‘intimately co-existing’ species), which are engaged in reciprocal selection. In specialist antagonistic interactions, or obligate mutualisms, each party exerts such strong selection on the other that there can be little differentiation between populations (e.g. et al., 2004; Johnson et al., 2010; Lively & Dybdahl, 2000). A key challenge is to understand the ecological factors that generate such geographical mosaics of co-evolution and co-adaptation (Thompson, 2013).

Environmental differences among populations are known to cause variation in the outcome of any species interaction. These might arise from variation in the availability of an essential resource (e.g. Johnson et al., 2010), or the presence of a common enemy species (e.g. Hopkins et al., 2017; Sun & Kilner, 2020), or differences in the abiotic environment (e.g.Kersch & Fonseca, 2005). Theoretical analyses suggest that this variation, in turn, can cause dramatic differences in the trajectory of local co-evolution (e.g. Nuismer et al., 2000). Therefore, determining ecological correlates of variation in the extent of reciprocal selection between the same two partner species can potentially explain why there is geographical variation in the pattern of local adaptation among populations (e.g. Garrido et al., 2012; Gorter et al., 2016; Johnson et al., 2010).

However, in reality, interactions between species rarely involve just two partner species. For example, one insect species commonly pollinates more than one plant species, while each plant species can be pollinated by more than one insect species (e.g. J. N. Thompson, 2013). Similarly, one cactus species can be in a protective mutualism with multiple species of ants (Ness et al., 2006), while multiple bumblebee species *Bombus* spp. commonly interact with multiple mite species (Haas et al., 2019). Likewise, host species richness and abundance are positively correlated with parasite species richness (e.g. Hechinger & Lafferty, 2005). Multispecies associations are likely to generate variation in the strength of reciprocal selection. For example, multiple infection of different parasite species or strains can differentially influence the fitness among different host species interacting in the same community, depending on the susceptibility and tolerance of each host species (Friesen et al., 2017). Therefore understanding interactions among different strains or species of symbionts is likely to help account for geographical variation in the pattern of local co-adaptation between symbionts and their hosts.

Here we apply this rationale to understanding the extent of local co-adaptation between burying beetles *Nicrophorus vespilloides* (Coleoptera: Silphidae) and the species complex of *Poecilochirus carabi* mites with whom they interact (Mesostigmata: Parasitidae). Both the burying beetle and the mite require small vertebrate carrion to breed, but only the beetle is capable of flight. The mites travel with the beetle between breeding opportunities whilst sexually immature deutonymphs, attached as benign passengers to the adult beetle’s thorax. When they locate a dead body, the adult beetles convert the carrion into an edible nest for their young, and then stay with their brood to defend and nourish them. Meanwhile, the mites disembark, moult into the adult stage, mate, and produce offspring alongside the beetle larvae (Schwarz & Müller, 1992). Breeding alongside mites can be costly for beetles (De Gasperin et al., 2015; De Gasperin & Kilner, 2015; Nehring et al., 2019), which select adaptations in the beetles to mitigate these costs. When their duties of care are complete, adult beetles fly off in search of new reproductive opportunities, carrying with them the next generation of mites. The timing of the mite’s lifecycle is therefore closely aligned with the duration of beetle parental care (Schwarz & Müller, 1992).

*P. carabi* exists as a species complex because it comprises distinct lineages of mites that are each specialised to breed on different species of burying beetle *Nicrophorus* spp (Nehring et al., 2017). Each burying beetle species differs slightly in its duration of care. This has favoured divergent adaptation in the mite populations associated with each species of burying beetle, which in turn has generated distinct mite lineages (Canitz et al., 2021; Müller & Schwarz, 1990; Wilson, 1982). Nevertheless, mite lineages can still interbreed and cannot be told apart morphologically. The key way to distinguish them is behaviourally, through their relative preference for different burying beetle species (Brown & Wilson, 1992; Schwarz, 1996; Wilson, 1982). Previous work suggests that when sympatric *Nicrophorus* species do not differ much in their duration of care, then each species of burying beetles carries a mixture of the different mite lineages associated with each of the sympatric beetles and the mites hybridise across lineages (Brown & Wilson, 1992). However, in populations where *Nicrophorus* species differ considerably in their duration of care (Brown & Wilson, 1992; Scott, 1998), especially when larger beetle species have relatively longer larval development time than smaller beetle species, each species is more likely to carry its own specialist mite lineage and each mite lineage is more likely to be reproductively isolated (Brown & Wilson, 1992).

We focused on *N. vespilloides* and *P. carabi* inhabiting two neighbouring yet geographically isolated populations (Gamlingay and Waresley Woods) in Cambridgeshire, UK. Previous studies have shown that there are differences in the *Nicrophorus* guild inhabiting each woodland (Sun et al., 2020). Both woods contain the smallest UK burying beetle *N. vespilloides* and the larger burying beetle *N. humator*. However, only Gamlingay Woods is additionally routinely inhabited by the intermediate-sized *N. interruptus* and *N. investigator*. We tested the following predictions that:

1. *N. vespilloides* from Gamlingay Woods should be more likely to carry a mixture of mite lineages, whereas *N. vespilloides* from Waresley Woods should be more likely to carry the *N. vespilloides* lineage of mites, reflecting the different species that routinely co-exist in each wood. To test this prediction, we conducted two consecutive choice experiments for mites to choose among all four *Nicrophorus* beetles in the first generation of *P. carabi*, and then bred these mites based on their choice. Their offspring were tested again to evaluate the consistency in choice and the extent to which each mite lineage was mixed in each woodland.
2. *N. vespilloides* should have lower reproductive success when breeding alongside a mixture of *P. carabi* lineages than when breeding with its specialist *P. carabi* lineage, with whom the beetle is more likely to be co-adapted. We experimentally manipulated the mite composition, either mixing all four lineages of *P. carabi* or allowing the pure *N. vespilloides* lineage of *P. carabi* to breed alongside *N. vespilloides*. At dispersal, we determined the reproductive success of both beetles and mites.
3. *N.vespilloides* and *P. carabi* should be divergently co-adapted in Gamlingay and Waresley woods, if they differ in whether they carry pure versus mixed lineages of mites. Specifically, we predicted that *N. vespilloides* and its specialist lineage of mites should be locally adapted to each other. We tested this prediction using reciprocal transplant experiments by exposing mites from either Gamlingay Woods or Waresley Woods to *N. vespilloides* beetles from Gamlingay and Waresley Woods.

Just as before, we determined the reproductive success of both beetles and mites at dispersal.

## Methods Study species

Gamlingay and Waresley Woods are fragments of the Wild Wood that covered England until deforestation (c. 1000-4000 years ago), and are roughly 2.5km apart (Sun et al., 2020). *N. vespilloides* are the most abundant burying beetles in each wood (81.6% in Gamlingay Woods and 93.7% in Waresley Woods (Sun et al., 2020)), and the *P. carabi* mite species complex is the most commonly found mite species associating with *Nicrophorus* beetles (Schwarz et al., 1998).

### Field observations

Surveys of burying beetle communities were conducted in Gamlingay (Latitude: 52.15555°; Longitude: −0.19286°) and Waresley (Latitude: 52.17487°; Longitude: −0.17354°) Woods in Cambridgeshire, UK. From June to October in 2016-2017, five traps at each site were baited with mouse carcasses and hung in the same location, separated by at least 150 m. We checked the traps every 2-3 weeks and collected all *Nicrophorus* spp. under permit. Traps were then refilled with fresh mice at each collection. The number and sex of each beetle species were recorded at each location for each trapping event. Each beetle’s pronotum width was measured to the nearest 0.01 mm as a standardised measurement of body size (Jarrett et al., 2017). We separated mites from their beetle hosts under anaesthetic by CO_2_. The data presented here are related to those from an earlier published study within this system that focussed only on the correlation between body size and mite load in *N. vespilloides* in 2017 (Sun et al., 2019). In the current study, we have used a more extensive dataset to thoroughly investigate the mite load on each *Nicrophorus* spp. during 2016-2017.

### Origin and maintenance of burying beetles and mites

Both species were kept under laboratory conditions at 21 ± 2°C and on a 16:8 light to dark cycle.

Beetles were kept individually in plastic boxes (12cm x 8cm x 2cm) filled with moist soil. Field-collected beetles were kept for at least two weeks before they were subjected to experimentation to even out any differences in sexual maturity and nutritional status. To breed beetles, *N. vespilloides* collected from the field sites were paired on a mouse in a breeding box lined with damp soil. All breeding boxes were then placed into cupboards to mimic underground environments. After eight days, we collected the dispersing larvae and transferred them to eclosion boxes (10 x 10 x 2 cm, 25 compartments) filled with moist soil. At eclosion, each emerging beetle was moved to a plastic container (12 x 8 x 2 cm) with moist soil. We fed beetles twice a week with minced beef for 2-3 weeks until they were sexually mature.

Mites were maintained in distinct populations, according to their woodland of origin, and kept apart from burying beetles except when breeding. To breed mites, each month we transferred 15 mite deutonymphs chosen at random, and a pair of beetles from the same population, to a new breeding box (17 x 12 x 6 cm with 2 cm of soil) furnished with a fresh mouse carcass (*n* = 10 for each population). After breeding, beetle parents and third-instar larvae were removed from the box. The mites remained and were given another adult beetle, and thereafter supplied with minced beef twice a week.

#### Prediction 1: Gamlingay N. vespilloides carry a mixture of P. carabi lineages

To investigate population differences in the number of mite lineages present in each wood, we used consecutive choice experiments. *P. carabi* from either Gamlingay or Waresley Woods were allowed to choose between one of the following field-collected burying beetle species: *N. vespilloides*, *N. humator*, *N. interruptus*, and *N. investigator*. Each species was represented by one individual, drawn at random from a pool of field beetles (184 *N. vespilloides*, 98 *N. humator*, 129 *N. interruptus* and 100 *N. investigator*). Burying beetles in this pool were haphazardly chosen from four field populations (Gamlingay Woods, Waresley Woods, Madingley Woods (Latitude: 52.22658°; Longitude: 0.04303°), and Thetford Forest (Latitude: 52.41286°; Longitude: 0.75167°). This allowed us to remove any beetle population-level effect on mite preferences, and to focus entirely on the effect of the beetle’s species on influencing mite preferences.

We used mites that had been bred for one generation in the lab so that the tested mites had no prior experience which might influence their choice of beetle. Four burying beetles (of the same sex, one from each species; *n* = 43 trials where beetles were all males and 38 trials where beetles were all females) were introduced into a plastic container (17 x 12 x 6 cm), around which they could move freely. At the same time 50 mites were also introduced – either from the Gamlingay population or from the Waresley population. It is unlikely that the mites hindered each other in attaching to host beetles because wild-caught beetles can carry up to 200-300 mites individually, even on *N. vespilloides*, the smallest of the UK species (Schwarz et al., 1998; Sun et al., 2019). The container held moist soil to a depth of 2 cm and minced beef in *ad libitum* quantities. All beetles and mites were starved for a day prior to the experiment to increase the likelihood that they gathered together on the mince, to mimic the natural context in which mites transfer between beetles hosts at feeding/ breeding opportunities. The number of mites carried by each beetle species was recorded 24 h later, and used to assess the mixing of different mite lineages in each population.

Mites that chose *N. vespilloides*, *N. humator*, *N. interruptus*, and *N. investigator* were defined as P-ves, P-hum, P-int, and P-inv, respectively. We then bred these mites separately on a fresh mouse carcass, with one mouse carcass for each lineage of mite identified by the choice test. The offspring of these breedings were then tested again for their burying beetle preferences, as a further test of extent to which the mite lineages were mixed in each woodland. For this second choice test, we introduced 10 mites and one beetle from each of the four species of the same sex (*n* = 92 trials where beetles were all males and 50 trials where beetles were all females) in a plastic container, and counted the number of mites on each beetle after 24 h. We tested 10 mites in the second choice experiment because we harvested relatively few P-hum and P-inv in the first choice experiment, and this meant there were fewer offspring available for testing.

#### Prediction 2: A mixture of P. carabi lineages negatively affects

N. vespilloides These experiments were focused on *N. vespilloides* and *P. carabi* mites drawn from Gamlingay Woods. We experimentally manipulated the composition of the mite community (= a group of 10 deutonymphs) associated with each burying beetle, generating two treatments: a) pure *N. vespilloides* lineage of *P. carabi* and b) a mixture of all four lineages of *P. carabi*. In the mixture, we introduced 3 P-ves, 1 P-hum, 4 P-int, and 2 P-inv, based on the relative proportion of the four *P. carabi* lineages from *Prediction 1*. The mites used were descendants of the second generation of *P. carabi* from the experiment above, and lineages were determined from the preferences they exhibited in the second generation of choice experiment. Ten mites were introduced at beetle pairing, directly onto the carcass. Pairs of beetles were sequentially assigned to one of the three mite treatments, introduced into a breeding box (17 x 12 x 6 cm with 2 cm of soil) and given a 15-20 g (17.71 ± 0.16) mouse carcass to breed upon. At larval dispersal, i.e. 8 days after pairing, we counted all larvae and weighed the whole brood (to the nearest 0.1mg). We calculated the average larval mass for each brood (total brood mass divided by the number of larvae). To determine the reproductive success of mites, we used CO_2_ to detach dispersing mite deutonymphs from adult beetles, at the end of the breeding event.

#### Prediction 3: Populations have adapted divergently

To assess the extent of local adaptation, we adopted a fully factorial design of experimental reciprocal mite infestation (e.g. Blanquart et al., 2013; Garrido et al., 2012; Nuismer & Gandon, 2008). These experiments were carried out in three blocks. Each beetle population (Gamlingay/Waresley) was infested with either 10 Gamlingay mite deutonymphs or 10 Waresley mite deutonymphs. Mites were introduced directly onto 15-20 g carcasses (16.97 ± 0.12 g), when beetles were paired. Pairs of beetles were unrelated, to prevent inbreeding. We took the same measurements of beetle and mite reproductive success as in the previous experiments, when larvae dispersed away from the carcass 8 days after pairing.

### Statistical analyses

We analysed the data using generalized linear mixed models (GLMM) with the glmer function in the *lme4* package in R version 3.4.3 (R Development Core Team, 2014). To obtain minimal adequate models, we applied a stepwise approach to exclude non-significant variables and interactions (Crawley, 2007). We included block as a random effect in all models, where appropriate. Tukey HSD tests were used for *post-hoc* pairwise comparisons, as necessary, using the *lsmeans* package (Lenth, 2016). All data are provided in electronic supplementary material.

#### Prediction 1: Gamlingay N. vespilloides carry a mixture of P. carabi lineages

We analysed variation in the number of mites carried by each beetle species using a Poisson model, with an offset of the log total number of mites allowed to make a choice in each trial. We included mite population (Gamlingay/Waresley), beetle species (*N. vespilloides*, *N. humator*, *N. interruptus*, *and N. investigator*), and their interaction as explanatory variables. Sex and body size of beetles were included as covariates. We also included as random factors the sampling year (2017 or 2018), and trial ID. We analysed the data in a similar manner for the second choice experiment, testing for consistency of beetle species preference among mites in the next generation. Of the 16 possible outcomes this two generation experiment yielded (N = 4 species for mites to choose in Gen 1 x N = 4 species for mites to choose in Gen 2), we analysed only those the mites that made the same choice of beetle species as their parents (i.e. N = 4 outcomes).

#### Prediction 2: A mixture of P. carabi lineages negatively affects

N. vespilloides We used generalized linear models (GLM) to analyse three measures of beetle reproductive success when exposed to different experimental combinations of mite lineages: brood size (using a Poisson distribution), brood mass and average larval mass (both using a Gaussian distribution). We included mite treatment as an explanatory variable, and additionally included carcass mass as a covariate when analysing brood size and brood mass. For analysis of average larval mass, we included larval density (brood size divided by carcass mass) as a covariate. We also analysed variation in mite reproductive success using a negative binomial GLM with mite treatments and carcass mass as explanatory variables.

#### Prediction 3: Populations have adapted divergently Local adaptation of beetles to mites

We analysed three measures of beetle reproductive success when exposed to different mite populations using GLMMs: brood size, brood mass and average larval mass. Beetle wood of origin (Gamlingay/Waresley), mite wood of origin (Gamlingay/Waresley) and their interaction were included as explanatory variables. We included carcass mass as a covariate for the analysis of brood size and brood mass, whereas we included larval density (brood size divided by carcass mass) as a covariate for the analysis of average larval mass. In all models, block was included as a random factor.

#### Local adaptation of mites to beetles

We analysed the number of dispersing mites deutonymphs present at the end of each reproductive bout using a negative binomial GLMM. Beetle wood of origin (Gamlingay/Waresley), mite wood of origin (Gamlingay/Waresley), and their interaction were included as explanatory variables. Carcass mass was included as a covariate, whereas block was included as a random factor.

## Results

### Field observation

In total, 1465 *Nicrophorus* individuals were caught over the two sampling years (780 and 685 for Gamlingay and Waresley Woods, respectively), carrying a total of 17,249 *P. carabi* mite deutonymphs on four beetle species (Fig. 1). We found that the mite load on each *Nicrophorus* species varied differently between populations (beetle species x population interaction, χ*²* = 46.19, d.f. = 3, *p* < 0.001; Fig. 1). Specifically, *N. vespilloides* from Gamlingay had an average lower number of mites compared to *N. interruptus* (*post-hoc* comparison: *z* = -2.86, *p* = 0.022), but carried similar number of mites compared to those of *N. humator* (*post-hoc* comparison: *z* = -0.73, *p* = 0.886) and *N. investigator* (*post-hoc* comparison: *z* = -1.53, *p* = 0.418). In Waresley Wood, however, *N. vespilloides* carried more mites than *N. humator* (*post-hoc* comparison*: z* = 6.80, *p* < 0.001), but we could detect no difference in the mite load carried by *N. vespilloides* and *N. interruptus* (*post-hoc* comparison: *z* = 2.22, *p* = 0.118), nor between *N. vespilloides* and *N. investigator* (*post-hoc* comparison: *z* = 1.14, *p* = 0.663). Moreover, comparing mite abundance on the beetle species, between woodlands, we found that Gamlingay *N. vespilloides* had lower number of mites than Waresley *N. vespilloides* (*post-hoc* comparison: *z* = -4.18, *p* < 0.001). In contrast, *N. humator* from Gamlingay had higher mite abundance than *N. humator* from Waresley (*post-hoc* comparison: *z* = 4.45, *p* < 0.001), and there was a tendency for Gamlingay *N. interruptus* to carry more mites than those from Waresley (*post-hoc* comparison: *z* = 1.72, *p* = 0.086). We could detect no significant difference between Gamlingay and Waresley in mite abundance on *N. investigator* (*post-hoc* comparison: *z* = 0.90, *p* = 0.370).

**Figure 1.**
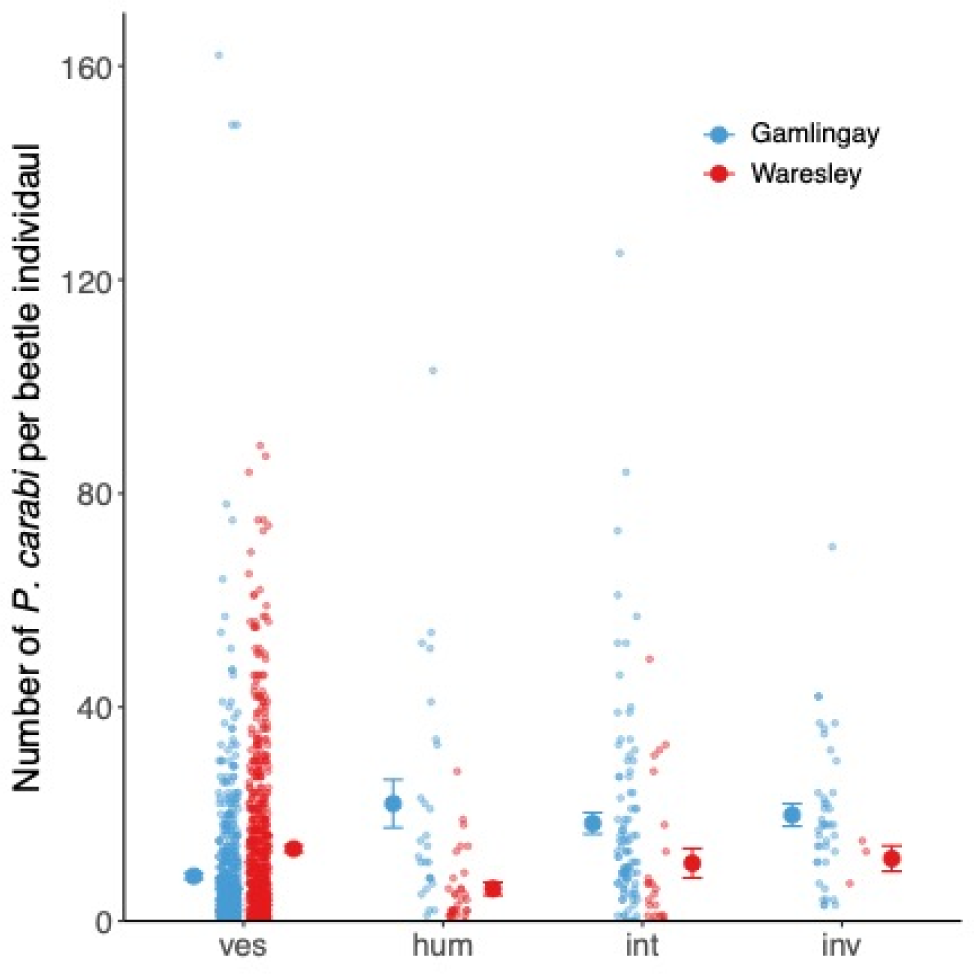
Number of *P. carabi* carried by each *Nicrophorus spp.* in Gamlingay and Waresley Woods. *N. vespilloides*, *N. humator*, *N. interruptus*, and *N. investigator* were represented as ves, hum, int, and inv, respectively. Means with standard error are shown, whereas individual data were pointed with a jitter effect to prevent overlap. (Note that *N. interruptus* and *N. investigator* are routinely found in Gamlingay Woods, though not in Waresley Woods – see Sun et al. 2020 for further details)

### Prediction 1: Gamlingay *N. vespilloides* carry a mixture of *P. carabi* lineages

The first lab-bred generation of *P. carabi* differed significantly in their relative preference for the different *Nicrophorus* species, depending on whether they were derived originally from Gamlingay or Waresley Woods (mite population x beetle species interaction, χ*²* = 152.62, d.f. = 3, *p* < 0.001; Fig. 2a; Table 1). *P. carabi* derived from Gamlingay Woods and Waresley Woods showed divergent preferences for the different *Nicrophorus* species (Gamlingay: χ*²* = 294.08, d.f. = 3, *p* < 0.001; Waresley: χ*²* = 459.69, d.f. = 3, *p* < 0.001). Gamlingay mites were similarly likely to favour *N. interruptus* and *N. vespilloides*. They favoured *N. investigator* less frequently and *N. humator* even less (Fig. 2a; Table 1). In contrast, Waresley *P. carabi* showed a clear preference for *N. vespilloides*. Their next-favoured beetle species was *N. interruptus*, followed by *N. investigator*, and then *N. humator* (Fig. 2a; Table 1). We used *post-hoc* comparisons to compare the strength of the mite preference for each *Nicrophorus* species between populations. We found that Gamlingay mites showed stronger preference for *N. humator* (*z* = 5.99, *p* < 0.001) and *N. interruptus* (*z* = 5.45, *p* < 0.001) than mites from Waresley. They were also slightly inclined to favour *N. investigator*, but this pattern was not statistically significant (*z* = 1.93, *p* = 0.054). By contrast, Waresley mites showed a higher preference for *N. vespilloides* than Gamlingay mites (*z* = -9.04, *p* < 0.001).

**Figure 2.**
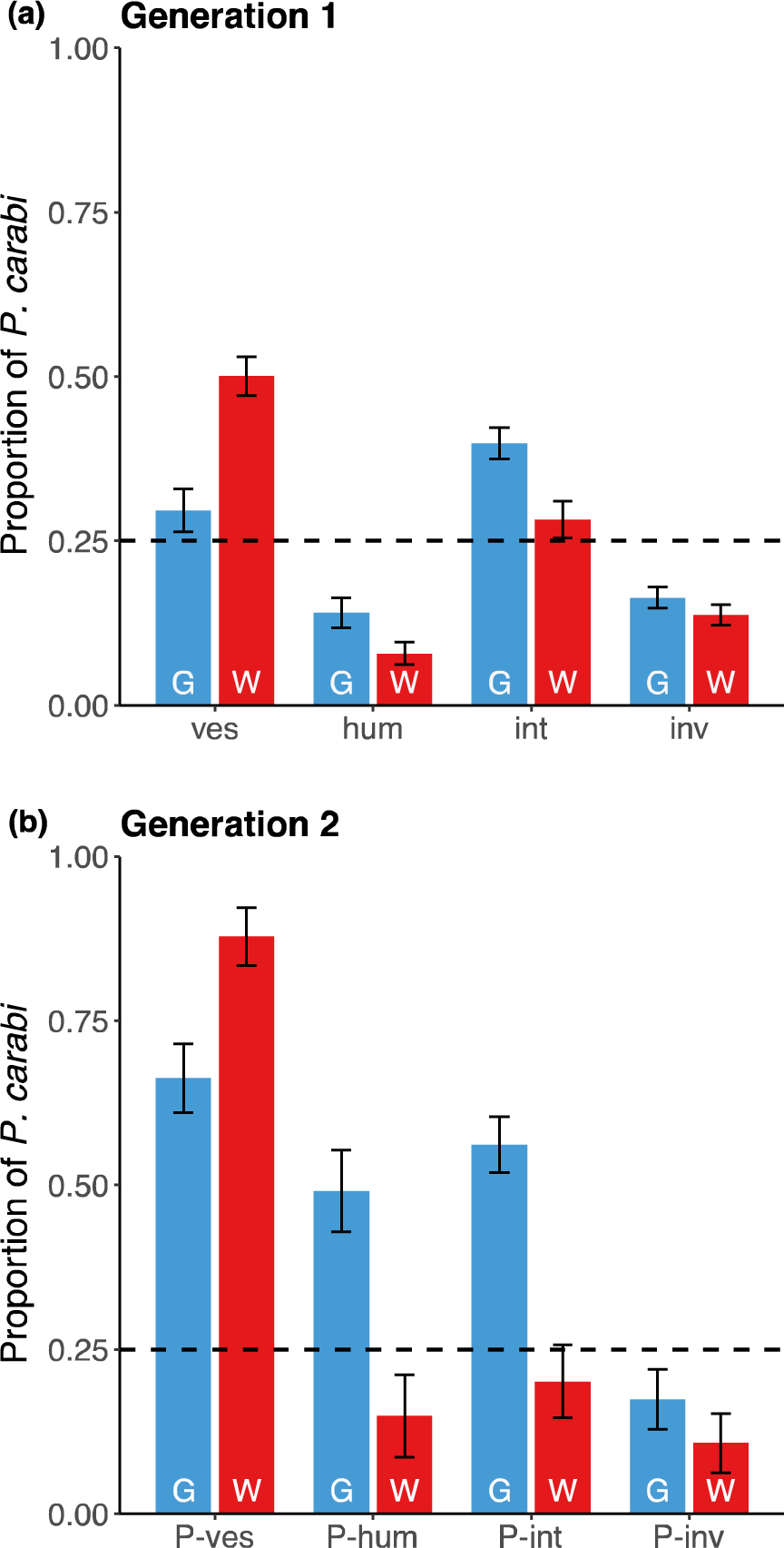
Population differences of mite preferences between Gamlingay (G) and Waresley (W) Woods, tested over two generations. (a) Proportion of mites that were attracted to each *Nicrophorus* spp. in the first generation (ves = *N. vespilloides*, hum = *N. humator*, int = *N. interruptus*, inv = *N. investigator*) and (b) proportion of mites that were attracted to their parents’ preferred *Nicrophorus* spp. P-ves, P-hum, P-int, and P-inv represent mites that chose *N. vespilloides*, *N. humator*, *N. interruptus*, and *N. investigator*, respectively. Means with standard error are shown. The dashed line at 25% represents the proportion of *P.carabi* associating simply by chance with one of the *Nicrophorus* species.

**Table 1.**
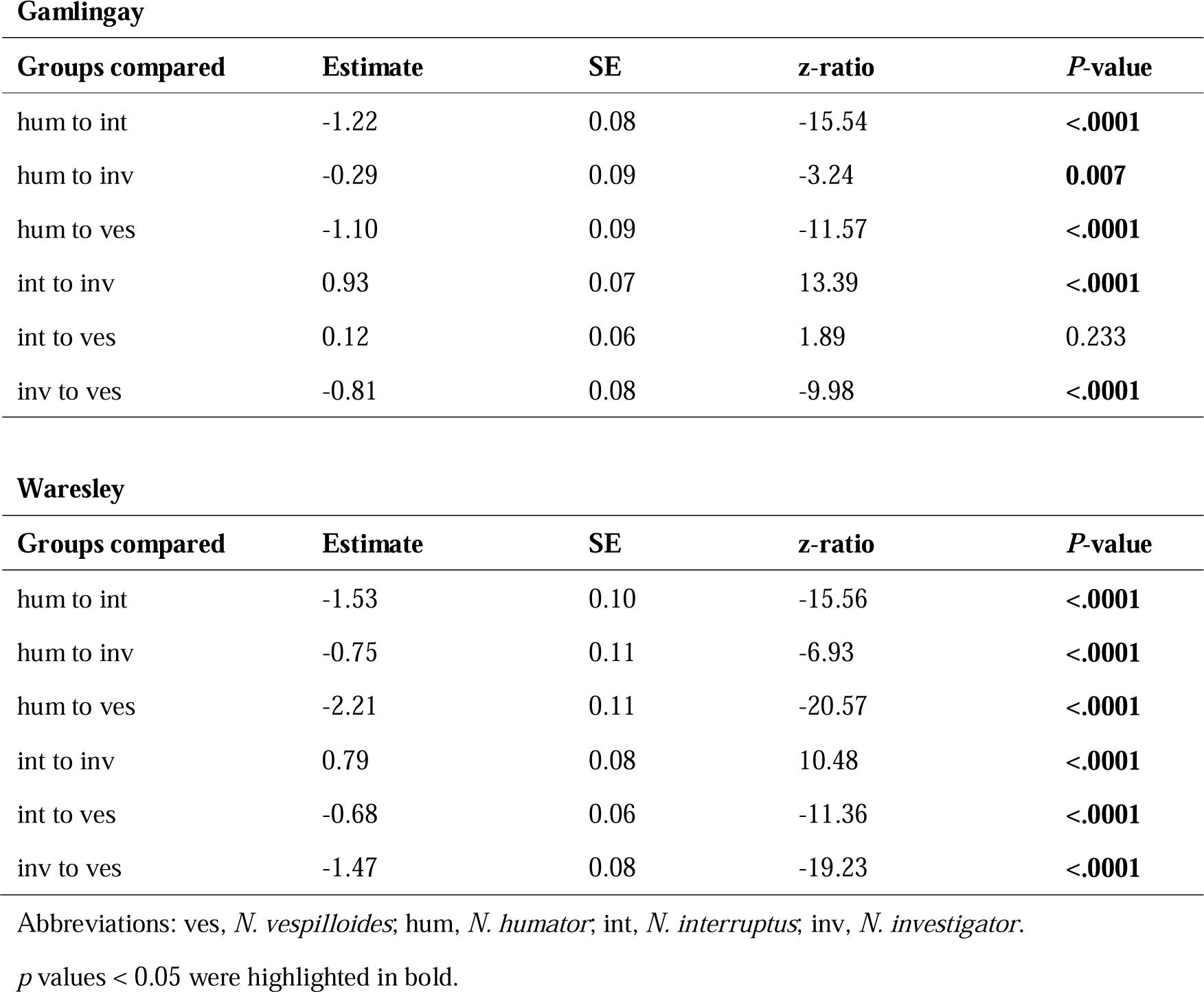
Results of Tukey’s post hoc comparisons for beetle species x population interaction in the first choice experiment.

We bred mites that showed the same preference for burying beetle species, and tested whether the preferences of the offspring matched those of their parents, to test for indirect evidence that mites were segregating into genetic lineages, as inferred from their host preference. The extent to which the beetle preferences aligned between the generations varied by woodland (mite population x beetle species interaction, χ*²* = 50.42, d.f. = 3, *p* < 0.001). We found that Waresley P-ves mites consistently had stronger preference for *N. vespilloides* than Gamlingay mites (*t* = -2.17, *p* = 0.030; Fig. 2b). In contrast, both P-hum and P-int mites from Gamlingay Woods showed stronger consistency in their choice of *N. humator* (*t* = 5.48, *p* < 0.001; Fig. 2b) and *N. interruptus* (*t* = 4.92, *p* < 0.001; Fig. 2b), respectively, compared to those from Waresley Woods. Gamlingay P-inv mites also shown a non-significant tendency to prefer *N. investigator* compared to Waresley P-inv mites (*t* = 1.78, *p* = 0.075; Fig. 2b).

### Prediction 2: A mixture of *P. carabi* lineages negatively affects *N. vespilloides*

We tested whether differential mixing of mite lineages in Gamlingay Woods caused variations in beetle reproductive success. We created experimental mite communities, manipulated to different degrees to contain mites with different beetle preferences. We found that beetles breeding alongside mites that varied in their preference for different beetle species produced smaller (χ*²* = 33.40, d.f. = 1, *p* < 0.001; Fig. 3a) and lighter broods (χ*²* = 5.39, d.f. = 1, *p* = 0.02; Fig. 3b) than beetles breeding with a pure population of *N. vespilloides-*specific mites. We found no effect of the experimental mite community composition on average larval mass (χ*²* = 2.55, d.f. = 1, *p* = 0.110; Fig. 3c). The beetle preferences of the mites also explained variation in mite reproductive success (Fig. 3d). Mites produced more offspring in a pure population of *N. vespilloides*-specific *P. carabi* than when in a mixture of different *P. carabi* ‘lineages’ (χ*²* = 18.13, d.f. = 1, *p* < 0.001).

**Figure 3.**
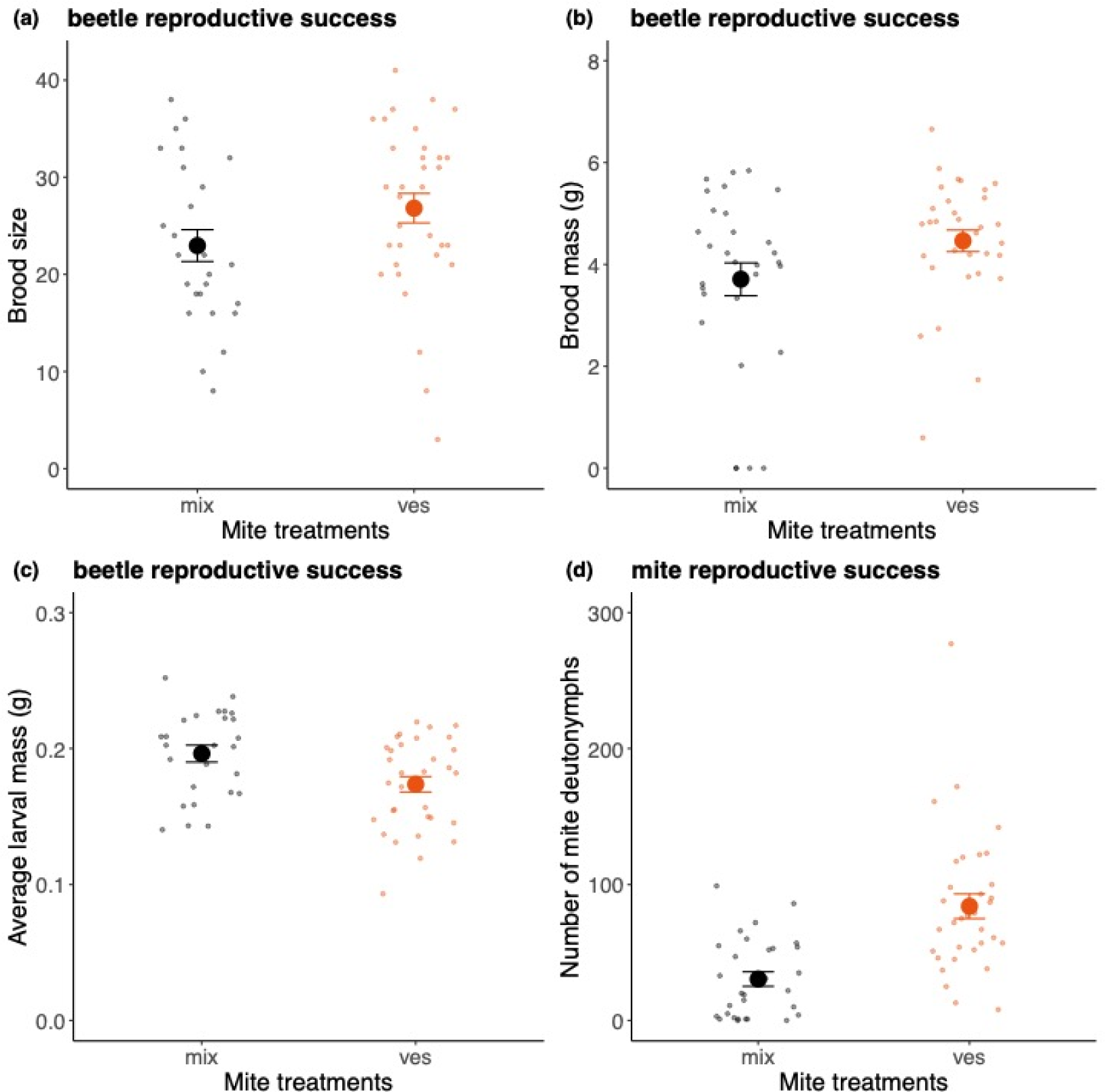
Reproductive success of beetles and mites from Gamlingay Woods, following experimental manipulations of the mite lineages on each carcass. Reproductive success of beetles was measured as (a) brood size, (b) brood mass and (c) average larval mass, whereas mite reproductive success was measured as (d) the number of deutonymphs dispersing with adult beetles. In the mite treatments, ‘mix’ means that beetles bred alongside 10 mites as a mixture of all four ‘lineages’ and ‘ves’ means that beetles bred alongside 10 mites from pure *N. vespilloides* lineage. Means with standard error are shown. Each point represents a different reproductive event.

### Prediction 3: Populations have adapted divergently

#### 1. Local adaptation of beetles to mite populations Brood size

Beetles exposed to Gamlingay and Waresley mites produced a similar brood size (Fig. 4a; Table 2). In addition, beetles from Waresley Woods consistently produced larger broods compared to those from Gamlingay Woods (Fig. 4a; Table 2).

**Figure 4.**
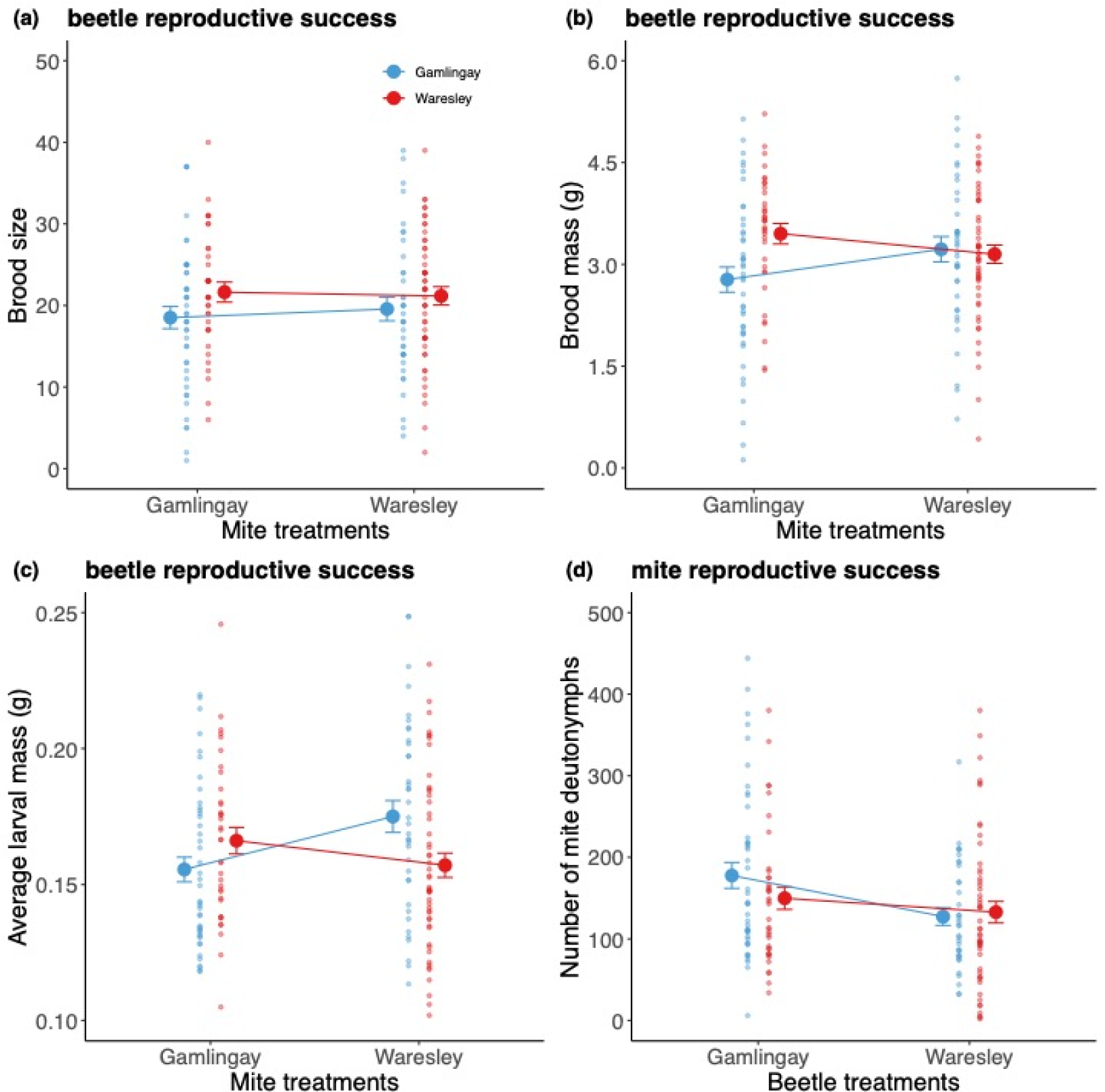
Burying beetle and mite reproductive success in relation to woodland of origin and mite/beetle treatments. Beetles / mites deriving originally from Gamlingay Woods are shown in blue, whereas those deriving originally from Waresley Woods are shown in red. The beetle / mite treatments refer to the origin of the partner species. Reproductive success of beetles was measured as (a) brood size, (b) brood mass and (c) average larval mass, whereas mite reproductive success was measured as (d) the number of deutonymphs dispersing with adult beetles. Means with standard error are shown. Each point represents a different reproductive event.

**Table 2.** Results from the final models analysing the fitness components of local adaptation between beetles and mites. *p* values < 0.05 were highlighted in bold.

| Dependent variable | Explanatory variables | $X^2$ | d.f. | *p* value |
| --- | --- | --- | --- | --- |
| Brood size | mite population | 0.67 | 1 | 0.412 |
|  | beetle population | 6.36 | 1 | **0.012** |
|  | carcass mass | 24.82 | 1 | **<0.001** |
| Brood mass | mite population | 5.19 | 1 | **0.023** |
|  | beetle population | 6.64 | 1 | **0.010** |
|  | carcass mass | 14.32 | 1 | **<0.001** |
|  | mite population*beetle population | 4.73 | 1 | **0.030** |
| Average larval mass | mite population | 16.69 | 1 | **<0.001** |
|  | beetle population | 13.38 | 1 | **<0.001** |
|  | larval density | 158.84 | 1 | **<0.001** |
|  | mite population*beetle population | 21.17 | 1 | **<0.001** |
| Number of mite offspring | mite population | 2.28 | 1 | 0.131 |
|  | beetle population | 7.75 | 1 | **0.005** |
|  | carcass mass | 0.40 | 1 | 0.526 |

*p* values < 0.05 were highlighted in bold.

##### Brood mass

We found that different mite populations affected burying beetle fitness in different ways, depending on the burying beetle’s woodland of origin (Table 2). In Gamlingay Woods, we found that Gamlingay beetles breeding with Gamlingay mites produced lighter broods compared to when they bred with Waresley mites (*post-hoc* comparison, local mites v. foreign mites: *t* = -2.28, *p* = 0.024). Beetles from Waresley Woods produced broods that did not differ significantly in mass whether they were breeding alongside Waresley mites or Gamlingay mites (*post-hoc* comparison, local mites v. foreign mites: *t* = 0.76, *p* = 0.451).

##### Average larval mass

We found that the woodland origin of the mites and of the beetles interacted in affecting average larval mass (Fig. 4c; Table 2). When beetles bred alongside mites from the same woodland population, their larvae were smaller than when they bred alongside mites from the other woodland for Gamlingay beetles (*post-hoc* comparison, local mites v. foreign mites: *t* = -4.08, *p* < 0.001) and for Waresley beetles (*post-hoc* comparison, local mites v. foreign mites: *t* = 2.35, *p* = 0.020), but the extent of reduction in average larval mass by breeding with local mites was greater in Gamlingay beetles than in Waresley beetles (Table 2).

#### 2. Local adaptation of mites to beetle populations

We found that the reproductive success of the mites depended on the woodland origin of the beetles but not that of the mites. Gamlingay mites had greater reproductive success when breeding alongside Gamlingay beetles rather than with Waresley beetles (Fig. 4d; Table 2). Gamlingay and Waresley mites performed equally well on Waresley beetles (Fig. 4d; Table 2).

## Discussion

The mosaic theory of co-evolution (Thompson, 2005) suggests that populations differ in the extent of co-adaptation between interacting species because the structure of selection varies between populations; because the strength of reciprocal selection varies between populations (Agrawal & Zhang, 2021); or because genetic variation influences the capacity for co-adaptation in different populations (Whitham et al., 2020); or some combination of all three of these factors. The novel contribution of this study is to show how interactions among the lineages of one species are themselves a source of selection, both within and between species (Sun et al., 2019) (Table 3).

**Table 3.**
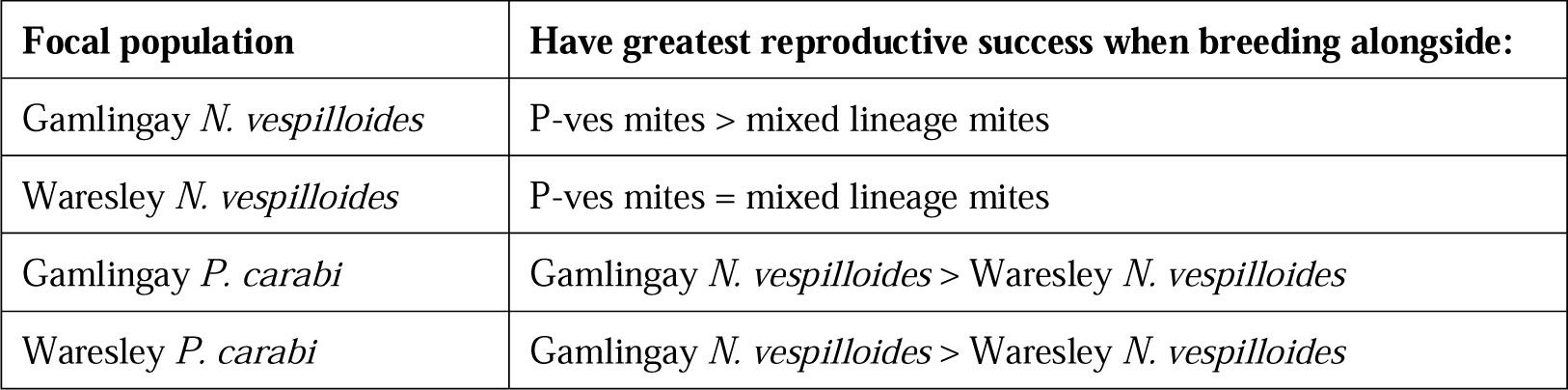
Summary of experimental results investigating the extent of local co-adaptation between *N. vespilloides* and *P. carabi* in Gamlingay *versus* Waresley Woods.

| Focal population | Have greatest reproductive success when breeding alongside: |
| --- | --- |
| Gamlingay *N. vespilloides* | P-ves mites > mixed lineage mites |
| Waresley *N. vespilloides* | P-ves mites = mixed lineage mites |
| Gamlingay *P. carabi* | Gamlingay *N. vespilloides* > Waresley *N. vespilloides* |
| Waresley *P. carabi* | Gamlingay *N. vespilloides* > Waresley *N. vespilloides* |

The differences we report here between Gamlingay and Waresley Woods can be traced back to the contrast in the *Nicrophorus* guild that inhabits each wood (Sun et al., 2020). In Gamlingay Woods, four *Nicrophorus* species co-exist. In support of Prediction (1), we found that *N. vespilloides* carries a mixture of mite lineages from all four *Nicrophorus* species. By contrast, in Waresley Woods, there are routinely only two burying beetle species (Sun et al., 2020). Waresley derived mites displayed a marked preference for associating with *N. vespilloides*, and this preference persisted between generations. Therefore, we conclude that Waresley *N. vespilloides* carries a near pure-bred lineage of P-ves mites. Perhaps *N. vespilloides* and *N. humator* differ too much in the duration of parental care for mixed lineages to persist in Waresley Woods (see Brown & Wilson 1992). This remains to be determined in future work.

Experimentally manipulating the lineages present in the mite community at each breeding event revealed two ways in which the composition of the mite community exerts selection. First, and unexpectedly, we found that mites themselves had lower breeding success when breeding as a mixed community than when breeding as a pure P-ves population. This suggests that the mite lineages impose selection on each other and that each mite lineage is locally adapted to breed alongside its own kind, perhaps through mechanisms that minimise local mate competition (Nehring & Müller, 2009). Male mites in particular are known to be lethally competitive (Nehring & Müller, 2009) and it would be interesting to determine whether the incidence of death is greater when the lineages are mixed.

Unexpectedly, we also found that *N. vespilloides* somehow modulates the selection that the mixed mite lineages impose upon each other during reproduction. Furthermore, the increase in mite reproductive success was more pronounced when Gamlingay *N. vespilloides* were present than when Waresley *N. vespilloides* bred alongside the mites, suggesting that the Gamlingay *P. carabi* mites have divergently adapted to breed in association with Gamlingay *N. vespilloides*.

From the Gamlingay *N. vespilloides*’ perspective, however, the increase in mite reproductive success came at the cost of the beetle’s reproductive success. Therefore the second way in which the mixed community of mites is divergent from the pure P-ves population lies in relative strength of selection it imposes on *N. vespilloides* – in support of Prediction (2). Perhaps as a direct consequence of increased competition among the mite lineages, or perhaps through other means, the mixed community more negatively affected Gamlingay burying beetle reproductive success than did the pure P-ves population of mites.

A key way in which mites reduce burying beetle reproductive success is by consuming beetle eggs and newly hatched larvae (de Gasperin & Kilner 2015, Sun & Kilner 2020 eLife). With greater numbers of mites breeding, potentially, in the mixed mite communities (Nehring & Muller 2009), perhaps there was a greater incidence of mite attack on beetle offspring. Consistent with this interpretation, we found that Waresley *N. vespilloides* actually had higher reproductive success than Gamlingay *N. vespilloides* when breeding alongside the mixed mite community. This could be because Waresley females lay more eggs than Gamlingay females (Sun et al., 2020) and so can better withstand the greater incidence of offspring attack by mites, as evidenced by Waresley beetles consistently producing larger broods, irrespective of mite presence and woodland of origin (Fig. 4a).

In summary, overall we find that *N. vespilloides* and the *N. vespilloides* mite lineage have co-adapted to each other (Table 3). However, neither *N. vespilloides* nor the *N. vespilloides* mite lineage is adapted to breed alongside other mite lineages. Local adaptation probably involves some form of tolerance rather than any specific defences against mite attacks on reproductive success (Svensson & Råberg, 2010). Since Gamlingay *N. vespilloides* carries a mixture of mite lineages, the extent of local adaptation between *N. vespilloides* beetles and their *P. carabi* mites is greater in Waresley Woods than in Gamlingay Woods.

Animals are commonly hosts to diverse communities of symbionts. The general conclusion we draw from our work is that variation in the extent of local adaptation between symbionts and their hosts depends on two factors that have been relatively overlooked: (1) the environmental effects that shape how these communities arise, and (2) the nature of interactions among different strains of symbiont. Nevertheless, each factor has been shown to be important in recent work on other species. For example, in deep-sea hydrothermal vent snails, environmental conditions have been found to have a greater effect than host genetic variation in determining symbiont composition and adaptation to local geochemical conditions (Breusing et al., 2022). Similarly, in pea aphids, cooperative relationships among symbiont strains have been found to influence the nature of host-symbiont interactions and adaptations (Peng et al., 2023). Whether these two factors are just as influential in influencing the extent of local adaptation between other species of hosts and their symbionts remains to be investigated.

## Acknowledgements

We thank the other members of the Kilner Group, especially S. Aspinall and C. Swannack for logistical support, and S. Issar for providing the field-caught beetles for the choice experiments.

## Data accessibility

The datasets supporting this article are available as electronic supplementary material and also have been deposited at Github (https://github.com/syuanjyunsun/local_adaptation).

## Authors’ contributions

S.-J. S. collected the data and performed the analyses. All authors designed the study and contributed to writing the manuscript.

